# High-yield fabrication of DNA and RNA scaffolds for single molecule force and torque spectroscopy experiments

**DOI:** 10.1101/661330

**Authors:** Flávia Stal Papini, Mona Seifert, David Dulin

## Abstract

Single molecule biophysics experiments have enabled the observation of biomolecules with a great deal of precision in space and time, e.g. nucleic acids mechanical properties and protein-nucleic acids interactions using force and torque spectroscopy techniques. The success of these experiments strongly depends on the capacity of the researcher to design and fabricate complex nucleic acid scaffolds, as the pertinence and the yield of the experiment strongly depend on the high quality and purity of the final scaffold. Though the molecular biology techniques involved are well known, the fabrication of nucleic acids scaffold for single molecule experiments still remains a difficult task. Here, we present new protocols to generate high quality coilable double-stranded DNA and RNA, as well as DNA and RNA hairpins with ~500-1000 bp long stems. Importantly, we present a new approach based on single-stranded DNA’s annealing and show, using magnetic tweezers, that it is more efficient to generate complex nucleic acid scaffolds in larger amount and at higher purity than a standard PCR-digestion-ligation approach. The protocols we describe here enable the design of any sort of complex nucleic acid scaffold for single molecule biophysics experiments and will therefore be extremely valuable to the community.

## Introduction

Genome expression and maintenance are at the heart of cellular activity, and are partly controlled by the mechanical properties of the genome, and the interactions of various enzymes with DNA and RNA. Single molecule force and torque spectroscopy techniques, e.g. stretch flow, acoustic-force spectroscopy, atomic force microscopy, optical tweezers and magnetic tweezers (1,2), have been very powerful to characterize the biomechanical properties of nucleic acids under force and torque (3–15), and nucleic acid-protein interactions (1). These studies were made possible by the successful design and fabrication of nucleic acid constructs, i.e. a scaffold of DNA or RNA that can be attached at one end to an anchor point and at the other end to a force transducer. For example, coilable DNA has been essential to study topoisomerases (16,17) and RNA polymerase transcription kinetics (18), whereas long stem nucleic acid hairpins, i.e. with a stem of 500-1000 bp, have been key to study either helicase or polymerase kinetics (19), and generally protein-nucleic acid interactions (20,21). The making of nucleic acid scaffolds for single molecule force spectroscopy experiments requires different standard biochemical reactions, e.g. polymerase chain reaction (PCR), in vitro transcription reaction (IVTR), digestion and ligation (17,22–28). Despite the apparent simplicity of these procedures, the inner nature of single molecule experiments adds an extra level of complexity, as it requires nucleic acid scaffolds with a high level of purity, and often results in a low yield due to the many intermediate purification steps. A standard approach to generate a DNA scaffold for a force and torque spectroscopy experiments consists of a PCR - restriction reaction - ligation reaction cycle (25). Complex DNA constructs, e.g. long stem DNA hairpins, may require several cycles of PCR-digestion-ligation to assemble into the designed scaffold, which eventually leads to a poor yield (21,22). The fabrication of RNA constructs differs significantly from that of DNA constructs, since RNA comes naturally single-stranded. Double-stranded RNA (dsRNA) constructs, either linear or with hairpins, are made by hybridization of single-stranded RNA (ssRNA) fragments obtained from IVTR, which may be ligated if required (26,29,30). However, coilable dsRNA and long stem RNA hairpins still remain a challenge to generate. Overall, new strategies to produce highly pure nucleic acid scaffolds at high yield are in a dire need from the single molecule community.

Here, we provide several detailed protocols using different strategies to reliably fabricate DNA and RNA scaffolds for single molecule force and torque spectroscopy experiments. Precisely, we developed a new strategy based on the hybridization of custom sequence of single-stranded DNA (ssDNA) to make long stem hairpins and linear coilable doublestranded DNA (dsDNA). We also provide new protocols to generate similar RNA scaffolds. Using high throughput magnetic tweezers (29,31,32), we evaluate quantitatively the quality and the purity of the scaffolds, and we show that our new strategy enables the fabrication of DNA hairpins in fewer steps and with a higher purity and yield than the existing methods (21,22). We also provide new protocols to synthesize long stem RNA hairpin, i.e. 500 bp, and linear coilable dsRNA with high purity and high yield. The different methodologies presented here constitute the basis for the fabrication of high quality nucleic acids scaffold for single molecule force and torque spectroscopy experiments and will therefore be of great interest to the community.

## Material and Methods

### DNA preparation

Plasmid or λ phage DNA was used as template for PCR (**Supplementary Table 1**). Biotin- and digoxygenin-labeled handles were amplified with a non-proofreading Taq DNA polymerase (*New England Biolabs (NEB), Ipswich, MA USA*.) and either biotin-16-dUTP or digoxigenin-11-dUTP (*Jena Biosciences GmbH, Jena, Germany*) was added to the PCR reactions to a final concentration of 40 μM, in addition to the 200 μM of nonlabeled dNTPs. Other DNA fragments were obtained by PCR with the Phusion High-Fidelity DNA polymerase (*Thermo Fisher Scientific, Waltham, MA USA*), except for the fragment containing the stem-loop (S-SL), which was obtained with the LA Taq DNA polymerase (*Takara Bio Europe*) using the GC buffer I. PCR products were purified using either the QIAquick PCR Purification Kit (*Qiagen, Hilden, Germany*) or the Wizard® SV Gel and PCR Clean-Up System (*Promega GmbH, Mannheim, Germany*). All primers were obtained from *biomers.net*. Unless otherwise stated, plasmids were obtained from the GeneArt Gene Synthesis Service (*Thermo Fisher Scientific*). Template DNA and primer design were performed with assistance of the molecular biology module of the Benchling platform.

### Restriction enzyme digest and ligation

Endonucleases were obtained from *New England Biolabs* (*NEB*) and reactions were performed following the manufacturer’s instructions. Restriction enzyme reactions were typically performed at 37°C for at least two hours in a final volume of 50 μl. T4 DNA ligase was used for DNA ligations overnight at 16°C in 1x T4 DNA ligation buffer. When necessary, final products were separated from non-ligated fragments by electrophoresis using a 0.8-1.5% (m/V) agarose gel and extracted from the gel with the Monarch® DNA Gel Extraction Kit (*NEB*).

Hybridization of complementary oligonucleotides or long overhangs was done in a thermocycler by heating the samples at 65°C for 15 minutes and then decreasing the temperature in 5°C steps every 5 minutes, until reaching 25°C. DNA products were then placed at 4°C or −20°C until further use.

For the DNA annealing strategy, PCR products obtained with *Taq* DNA polymerase were treated with T4 DNA polymerase (*NEB*) for 15 minutes at 12°C in 1x buffer 2.1 (*NEB*), to remove 3’-A overhangs.

### Nicking of the DNA and purification of ssDNA

Reactions with the nicking endonucleases Nb.BbvCI and Nt.BbvCI were set with 2 to 3 μg of DNA in 1x Cutsmart buffer, at 37°C for 2 hours. The samples were then purified with the Monarch® PCR & DNA Cleanup Kit (NEB) and eluted in 10 μl. 10 μl of denaturing RNA Gel Loading Dye (2X) (*Thermo Fisher Scientific*) was added to the samples and the mixture was incubated for 5 minutes at 75°C DNA. Single strands were separated by electrophoresis in a 1.2% (m/V) agarose gel stained with Sybr™ Gold (*Thermo Fisher Scientific*). After electrophoresis, the gel was left to rest in TAE buffer (40 mM Tris, 20 mM acetic acid and 1 mM EDTA, pH 8.5) for 30 minutes to remove excess staining dyes and reduce background signal.

If better resolution of single-stranded bands was desired, an alkaline agarose gel was prepared by dissolving agarose in boiling distilled water at 1.2% (m/V), and letting the agarose mix cool down to ~55-60°C before complementing it with an alkaline buffer, to a final concentration of 30 mM NaOH and 1 mM EDTA. One volume of 6X alkaline electrophoresis loading buffer (180 mM NaOH, 6 mM EDTA, 18% Ficoll 400, 0.05% bromcresol green) was added to 5 volumes of DNA and the samples were denatured for 5 minutes at 75°C before loading into the gel. The electrophoresis was carried out for approximately 4 hours at 3 V/cm in the same buffer. The gel was then neutralized for 30 minutes in 0.5 M Tris-HCl buffer, pH 7.5, stained in a 1X Sybr®Gold solution for 30 minutes and incubated in the neutralizing buffer for 30 minutes to reduce background staining.

The ssDNA fragments were identified using a blue light transilluminator (**Supplementary Figure 1**), cut out of the gel and purified using the Monarch® DNA Gel Extraction Kit. The concentration of the recovered DNA was measured by absorption using a Nanodrop (*Thermo Fisher Scientific*).

### ssDNA annealing and ligation

Annealing reactions were set in a total volume of 40 μl, in a final buffer concentration of 10 mM Tris pH8, 1 mM EDTA and 50 mM NaCl. Around 50-200 ng of each singlestranded DNA were added to the reaction, corresponding to 0.2 – 0.5 pmol of each fragment. Samples were heated to 65°C for one hour in a thermocycler and the temperature was slowly ramped down by 1.2°C/5 minutes, until it reached 25°C. Samples were purified with the Monarch® PCR & DNA Cleanup Kit. For torque experiments, nicks in the final DNA constructs were ligated with T4 DNA ligase at 16°C, in 1x T4 ligase buffer (*NEB*). It is crucial to purchase PCR primers that are phosphorylated at the 5’-end and to remove 3’-A overhangs left by Taq polymerase on the handles after PCR, otherwise the ligation of the nicks cannot take place.

### RNA preparation

Plasmid DNA (pBB10 or pMTM2, **Supplementary Table 1**) was used as template for PCR with primers containing the T7 promoter (**Supplementary Table 2**). Purified PCR products were used as template to transcribe RNA *in vitro*, using the RiboMAX™ Large Scale RNA Production System, T7 (Promega GmbH, Mannheim, Germany), as previously described (Dulin *et al*., 2015b). RNA was purified with the RNeasy MinElute kit (*Qiagen*) and concentrations were determined using a Nanodrop.

### RNA annealing

One picomole (pmol) of the BIO and DIG handles and 1.5 - 2 pmol of the other ssRNAs were used for the annealing reactions, which were set in a total volume of 100 μl, in 8.5 mM sodium citrate and 7.5 mM NaCl, pH 6.4. Annealing was done in a thermocycler as described for ssDNA samples. Double-stranded RNA was purified with the RNeasy MinElute cleanup kit (*Qiagen*). Hybridization products were eluted with 1 mM sodium citrate. Concentrations were determined using a Nanodrop.

### RNA ligation

For the assembly of torsionally constrained dsRNA constructs and for the RNA hairpin, five micrograms of each RNA sample were treated before annealing, first with Antarctic Phosphatase and subsequently with T4 Polinucleotide Kinase (*NEB*), following the manufacturer’s instructions, to obtain monophosphorilated 5’ ends needed for ligation. The annealed strands were ligated with T4 RNA ligase 2 (*NEB*), in 20 μl reactions for 1h at 37°C. RNA samples were purified after each of the steps above (dephosphorylation, phosphorylation and ligation) with the RNA Clean & Concentrator™-5 (Zymo Research Europe, Freiburg, Germany) and eluted with nuclease-free water. Long-term storage of RNA was done in The RNA Storage Solution (Invitrogen, Carlsbad, CA, USA).

### High throughput magnetic tweezers

To control the quality of the nucleic acid scaffolds, we used the high throughput magnetic tweezers apparatus previously described in Ref. (29,31,32). A typical field of view (FoV) is shown in **Supplementary Figure 2**. Shortly, it is a custom inverted microscope with a 50x oil immersion objective (CFI Plan Achro 50 XH, NA 0.9, *Nikon, Germany*), on top of which a flow chamber is mounted; magnetic beads are tethered to the bottom glass coverslip by the nucleic acid scaffold (**Figure 1**). To apply an attractive force to the magnetic beads and stretch the nucleic acid tether, a pair of vertically aligned permanent magnets (5 mm cubes, SuperMagnete, Switzerland) separated by a 1 mm gap are positioned above the objective (31); the vertical position and rotation of the beads are controlled by the M-126-PD1 and C-150 motors (*Physik Instrumente PI, GmbH & Co. KG, Karlsruhe, Germany*), respectively. The field of view (FoV) is illuminated through the magnets’ gap by a collimated LED-light source located above it, and is imaged onto a large chip CMOS camera (Dalsa Falcon2 FA-80-12M1H, Stemmer Imaging, Germany).

**Figure 1:**
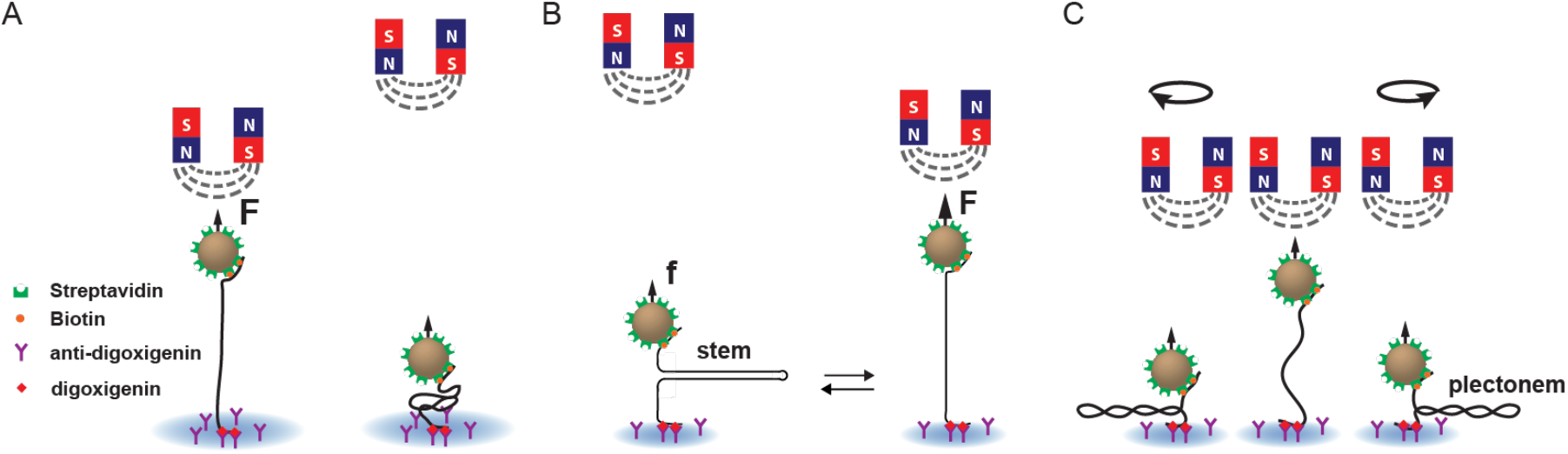
Force and torque spectroscopy experiments on nucleic acids using magnetic tweezers. To tether the magnetic bead to the surface of the flow chamber, the nucleic acid scaffold has two handles, one labeled with multiple biotins and the other with multiple digoxygenins. The biotins bind to multiple streptavidins that coates the magnetic bead, and the digoxygenins bind to multiple anti-digoxygenins coating the top surface of the bottom coverslip of the flow chamber. A pair of permanent magnets located above the flow chamber generates an attractive force (F, f) on the magnetic bead that stretches either **(A)** the linear or **(B)** the hairpin nucleic acid (NA) tether. **(C)** Rotation of the magnets causes rotation of the magnetic bead, which supercoils a torsionally constrained, i.e. coilable, dsNA tether, i.e. with multiple attachment points at each end and no nicks, and forms plectonems for both positive and negative supercoils at low force, e.g. ~0.3 pN.

### Flow Cell assembly and preparation

The flow cell assembly has been described previously (31). The flow cell was mounted on the magnetic tweezers setup and rinsed with 1 ml 1x Phosphate buffered saline (PBS). 100 μl of a 1:1000 dilution of 3 μm polystyrene beads (LB30, *Sigma Aldrich, Germany*) were added and after a 3-minute incubation, the flow cell was rinsed with 1 ml PBS. 40 μl of anti-digoxigenin Fab fragments (1 mg/ml) were added and the excess rinsed away with 1ml PBS after 30 minutes incubation. The flow cell was then treated with bovine serum albumin (BSA, *New England Biolabs*) (100 mg/ml) for 10 minutes and rinsed with 1 ml PBS. To remove BSA excess from the surface, high salt buffer (10 mM Tris, 1mM EDTA pH 8.0, supplemented with 750 mM NaCl and 2 mM sodium azide) was flushed through the flow cell and 10 minutes later washed away with measurement buffer (TE supplemented with 150 mM NaCl and 2 mM sodium azide). Nucleic acid constructs (~0.025 ng/μl) were mixed with either 10 μl of MyOne (Invitrogen) beads for the rotation-extension experiments or 20 μl of M270 beads (Invitrogen) for the force-extension experiments with hairpins, and were incubated in the flow cell for ~5 minutes before flushing away the non-tethered magnetic beads with measurement buffer.

### Magnetic tweezers measurement and analysis

The data were acquired in real-time at 58 Hz and the (x, y, z) trajectories of the tethered magnetic beads were drift-corrected using a reference bead attached to the glass surface of the flow chamber (32). We made a first selection of the tethers based on length, by measuring the difference in tether extension between ~0 pN and ~8 pN, and we selected only the tethers that had an extensions within ~10% of the expected crystallographic length. Coilable single nucleic acid tethers experiencing a 4 pN force have to demonstrate either a constant or a decrease extension in extension when they are either negatively or positively supercoiled, respectively (30,33), whereas non-coilable single nucleic acid tethers show no change in extension, and multiple nucleic acid tethers show a decrease in extension for both rotation directions. We report here the ratio of coilable tethers over the total number of tethers.

At ~0.3 pN, the extension of coilable tethers was measured at rotations between −20 and 20 turns (linear DNA) or −15 to 20 turns (linear RNA) in steps of 0.5 turns for 12 s per step. At ~4 pN, the turn range for linear RNA was increased from −15 to +35 turns. For the analysis, only the position at every full turn was evaluated. The mean extension of the tethers was calculated for the last six seconds at each magnet position measurement. The extension-rotation plots were centered to have the maximum tether extension at ~0.3 pN at the zero turn position.

For hairpins, the force was increased from ~0.1 pN to ~25 pN and subsequently decreased to ~0.1 pN at a constant magnet speed of ~0.4 mm/s. Tethers showing a jump-like change in extension of total amplitude of either ~0.35 μm, i.e. 500 bp stem, or ~0.7 μm, i.e. 1000 bp stem, were counted as hairpins.

### Magnetic tweezers data analysis

Each construct was made twice, except for the RNA hairpin and non-coilable linear RNA. For each tweezers measurement, the median of good tethers (coilable or hairpin) per FoV, the corresponding 95% confidence interval and the average number of good tethers/ng/FoV were obtained. From both replicates, the mean of the median and the mean error were extracted. Finally, the mean and the standard deviations for the number of good tethers/ng/FoV between the two tested replicates were calculated.

## Results

### Strategies to design double-stranded linear NA scaffolds

To generate linear coilable DNA scaffolds, we need a non-nicked stem flanked by handles being attached at multiple points to the beads at one end and to the surface at the other end (**Figure 1AC**). We use two different approaches to generate such DNA scaffold. The first approach has been extensively used and described in past reports (17,25), i.e. the linear DNA stem and the handles are obtained from PCR, followed by digestion and ligation reaction (**Figure 2A, Material and Methods**). We call this strategy ligated NA scaffold (LNS). We hypothesized that the LNS strategy bottleneck is the stability of the short hybridized region, i.e. ~2-6 bp, that supports the ligation reaction. Interestingly, dsRNA scaffolds fabrication shows a high conversion rate of the ssRNA’s into the final dsRNA (29), which likely originate from the long annealed region, i.e. hundreds of nucleotides, that increases the specificity and the stability of the scaffold. To verify whether long and complementary ssDNA molecules would increase the yield of high quality DNA scaffolds production, we need long complementary ssDNA’s with custom sequence. However, commercially available solid-phase synthesis DNA oligonucleotides are relatively short, i.e. up to ~200 nt, and expensive, while high throughput bacteriophage-derived strategy to generate long ssDNA, i.e. >200 nt, with a custom sequence still represents a challenge (34).

**Figure 2:**
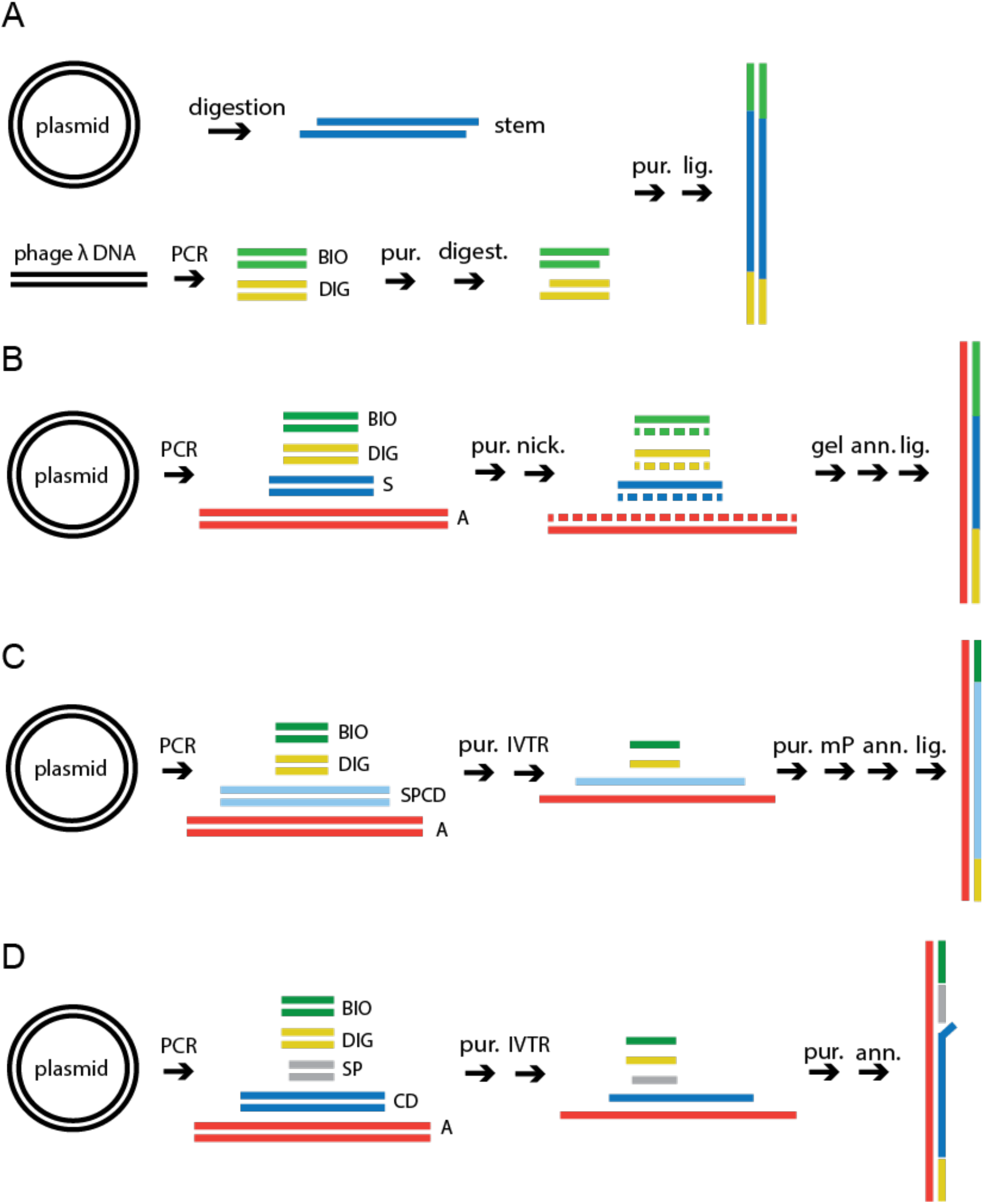
Experimental strategies to assemble linear DNA and RNA scaffolds. The colored lines represent different nucleic acid strands. BIO and DIG are respectively biotin- and digoxygenin-labeled **(A)** DNA construct, ligation method (LNS): linear or plasmid DNA is used as a template for restriction digestions and PCR reactions; fragments are purified (pur.) and ligated (lig.) to obtain the desired final product. **(B)** DNA construct, annealing method (ANS): plasmid DNA is used as template for PCR reactions; one strand is nicked and removed; complementary single strands are annealed (ann.) to obtain the desired final product. **(C)** RNA construct, coilable (ANS): RNA strands are obtained by run-off *in vitro* transcription reaction (IVTR), then purified and annealed. Single strands are monophosphorylated (mP) prior to annealing and then ligated (lig.) to obtain a coilable product. **(D)** RNA construct, non-coilable (ANS): template DNA is amplified by PCR and purified; RNA single strands are obtained as in (C) and annealed.

Therefore, we derived a new strategy using the engineered DNA nicking endonucleases to nick either the bottom strand, i.e. Nb.BbvCI, or the top strand, i.e. Nt.BbvCI (35), to generate at will ssDNA from dsDNA. These endonucleases have previously been used to introduce a modification in a DNA scaffold (23), e.g. a quantum dot, or to insert a small hairpin (36), but not to generate long ssDNA’s to synthesize either a full length coilable dsDNA or a long DNA hairpin based on annealing. To this end, we have designed a DNA plasmid with regularly interspaced insertions of BbvCI nicking sites (5’-CCTCAGC-3’). Using BbvCI nicking variants and the specifically designed plasmid, we are now able to generate ssDNA at will simply by performing PCR followed by digestion of either strand and gel electrophoresis purification. Furthermore, we chose to place the BbvCI nicking sites every 80 base pairs (bp) to clearly identify the non-nicked ssDNA from the nicked fragment during gel purification and extraction (**Supplementary Figure 1, Material and Methods**). The hundreds of nucleotides long ssDNA’s produced are subsequently annealed and ligated together to form a linear coilable dsDNA scaffold (**Figure 1C**, **Figure 2B**). We call this strategy annealed NA scaffold (ANS).

To generate a dsRNA scaffold using ANS, the fabrication is simpler than with DNA, as RNA comes naturally single-stranded from IVTR. However, if scaffold coilability is required, two intermediate reaction steps must be performed to convert the 5’-end triphosphate into a monophosphate and enable ligation of the strands (**Material and Methods**, **Supplementary Information**). To this end, the ssRNA’s are treated with the Antarctic Phosphatase and T4 Polynucleotide Kinase (NEB, Ipswich, MA). Thereafter, the ssRNA’s are annealed and ligated together to generate a coilable dsRNA (**Figure 2C**). It is important to mention that the ssRNA must anneal perfectly for the RNA ligation to work, i.e. without non-base paired nucleotides between them. Therefore, the PCR primers containing the T7 RNA promoter must be carefully designed, such as that the first base of the transcript will be the last G of the promoter sequence (TAATACGACTCACTATA**G**) (**Supplementary Figure 3**). Ideally, the annealing site of the primer in the template DNA should contain a GG sequence, since transcription is enhanced by one or two extra G’s at the transcription start site. When non-coilable dsRNA constructs are sufficient for experiments, e.g. for RNA-dependent RNA polymerase replication activity experiments (29,37,38), the ligation step may be skipped (**Figure 2D**).

### Generating NA hairpins using LNS and ANS

The LNS strategy to generate constructs containing long DNA hairpins has been previously described in detail (21,22). Shortly, multiple dsDNA fragments are amplified by PCR, digested and ligated together with other oligonucleotides to make a complete DNA hairpin (**Figure 3A, Material and Methods**). Given the relatively low efficiency of the ligation of multiple fragments, intermediate products have to be gel-purified after each ligation reaction and progressive loss of product takes place. As a consequence the protocol is time-consuming and the yield low. To circumvent these issues, we applied the ANS strategy to fabricate DNA hairpins (**Figure 3B**). We designed a plasmid that contains evenly spaced BbvCI nicking sites flanking a complementary inverted repeat, which will later form the hairpin by self-hybridization. Such DNA sequences represent a difficulty for standard PCR DNA polymerases, as high-fidelity DNA polymerases usually have poor strand displacement activity (39). Therefore, we used the LA Taq DNA polymerase (*Takara Bio* USA, Inc.) that passes efficiently through the complementary inverted repeat. Each DNA fragment is generated by PCR, followed by Nb.- or Nt.BbvCI nicking, and gel electrophoresis purification to extract the ssDNA’s that are annealed together (**Figure 3B, Material and Methods**). A final ligation reaction is not mandatory, since the annealing over hundreds of base pairs provides enough stability for the construct at forces below the overstretching force transition at ~50 pN (6,40). Advantageously, the ANS strategy requires fewer gel electrophoresis purification steps as compared to the LNS strategy (**Figure 3A**).

**Figure 3:**
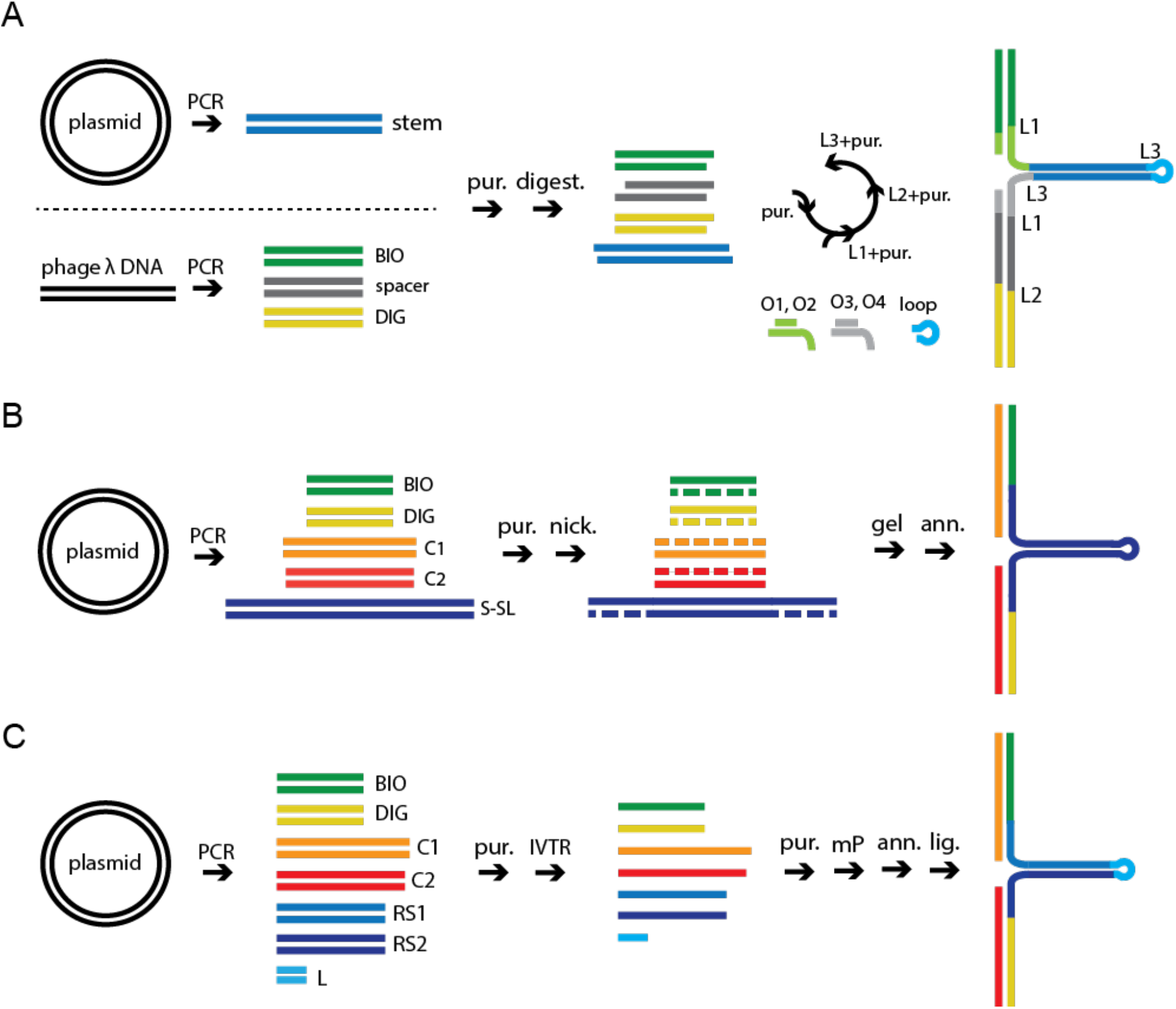
Experimental strategies to assemble long DNA and RNA hairpins. The colored lines represent different nucleic acid strands. BIO and DIG are respectively biotin- and digoxygenin-labeled. **(A)** DNA hairpin construct using LNS: linear or plasmid DNA is used as template for PCR reactions; amplified fragments are purified and digested; fragments are then submitted to three rounds of purification and ligation (L1, L2, L3) to obtain the desired final product. **(B)** DNA hairpin construct, annealing method (ANS): template DNA is amplified by PCR and purified (pur.); one strand of the amplified fragments is nicked with enzymes Nb.BbvCI or Nt.BbvCI, gel purified and annealed (ann.) to obtain the final construct. **(C)** RNA hairpin construct: template DNA is amplified by PCR and purified, stem is amplified in three separate parts; RNA products are obtained by IVTR, purified and monophosphorylated (mP); products are then annealed and ligated to obtained the final construct.

As previously mentioned, RNA comes naturally single-stranded from IVTR. However, we have not identified reaction conditions or an enzyme able to perform transcription through a long DNA inverted repeat without premature transcription termination. Therefore, the ANS strategy alone failed to produce an RNA hairpin with a long stem. We devised a strategy to assemble the hairpin stem in three parts that are subsequently joined in a final ligation step. To this end, we generated shorter, partially complementary ssRNAs that were annealed to form the long hairpin, and the resulting nicks in the dsRNA stem were ligated together (**Figure 3C**).

### LNS versus ANS to fabricate NA linear scaffolds

We tested the new constructs for quality and yield using the magnetic tweezers assay. The batch of linear nucleic acid construct produced is classified in three parts: the coilable designed construct, non-coilable designed construct – originating from improper ligation or nick(s) –, and junk DNA, i.e. the fraction in excess that did not assemble into a tethering scaffold. To assess the quality of the batch, we measured the percentage of coilable constructs (**Figure 4AB**). To assess the yield of the batch, we measured the fraction of coilable tethers obtained per nanogram (ng) of NA construct used in each experiment, i.e. per field of view (FoV). In short, the best strategy should produce the highest coilable fraction and the largest number of coilable tether per ng of DNA per field of view.

**Figure 4.**
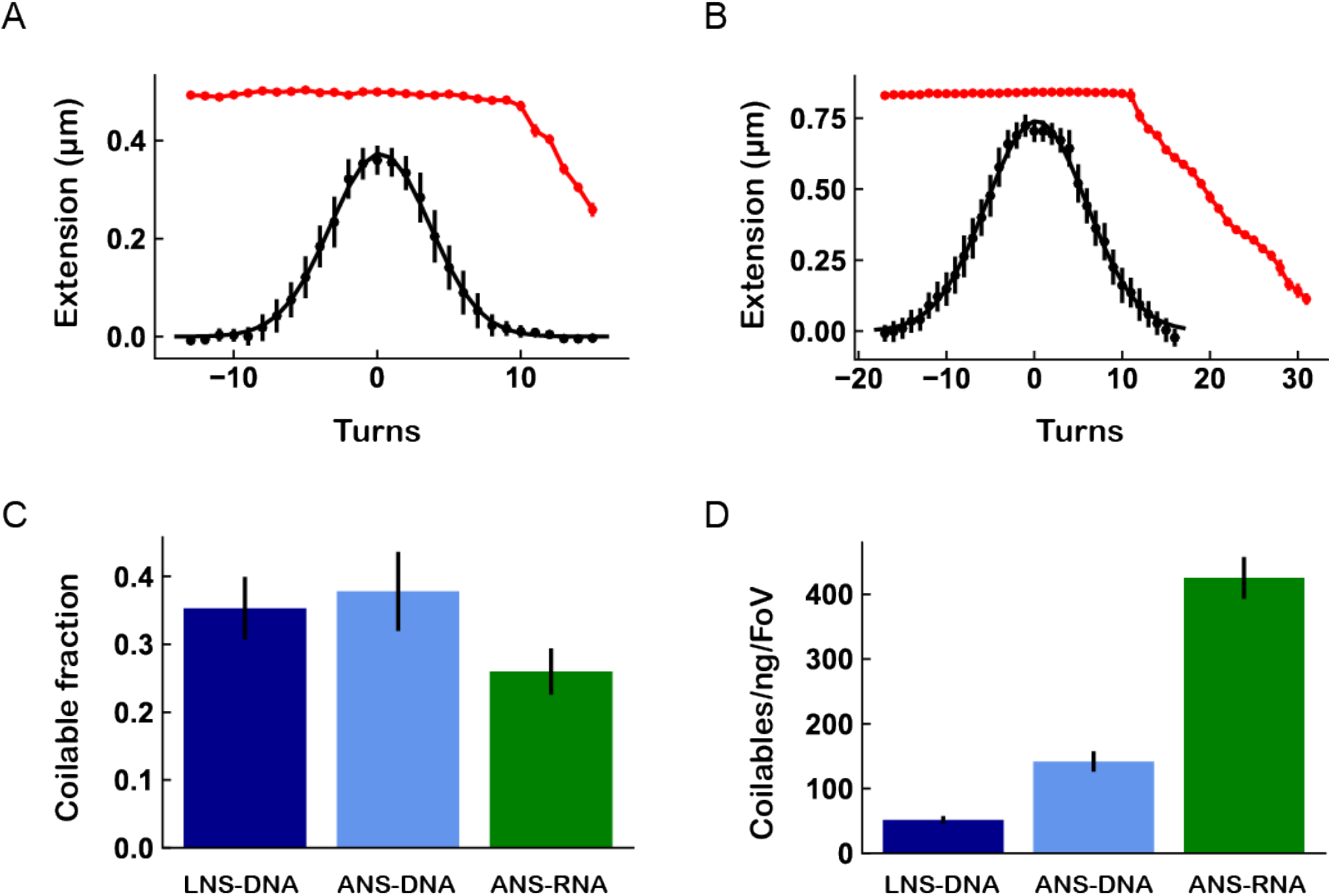
Comparative assessment of quality and yield for linear NS scaffolds using LNS and ANS methods. **(A,B)** Extension as a function of rotation for coilable 1.4 kbp dsDNA (A) and 4 kbp dsRNA (B), as described in **Figure 1C**. Z-positions of the beads were measured at 0.3 pN (black) and at 4 pN (red) forces. Data at 0.3 pN were fitted with a Gaussian function (solid black line). **(C)** Fraction of coilable tethers obtained for DNA or RNA constructs assembled using LNS or ANS strategies. **(D)** Scaffold recovery yield. The histogram shows the number of coilable tethers obtained per nanogram of nucleic acid scaffold per field of view.

To characterize the coilable fraction, we perform a rotation-extension experiment at two forces, 4 and 0.3 pN (**Figure 1C, Figure 4AB**). We measured the coilable fraction of tethers produced by LNS and ANS and observed that both methods provide a similar fraction of coilable molecules, i.e. 0.35 ± 0.05 and 0.36 ± 0.06 (median ± 95% confidence interval), respectively (**Figure 4C, Supplementary Table 3**). Although the mass of scaffold obtained was ~1.2-fold higher for the LNS strategy, 358 ng vs. 290 ng for ANS, the number of coilable tethers observed per field of view per ng of DNA used in one experiment was almost three times higher for the ANS than for the LNS batch, i.e. 142 and 51, respectively (**Figure 4D, Supplementary Table 3**). In other words, the ANS batch contained less junk DNA than the LNS batch, i.e. the fragments of DNA assemble more efficiently.

For the dsRNA scaffolds, we observed a coilable fraction of 0.26 ± 0.03 (**Figure 4C**). Previously, dsRNAs have been produced using a stalled RNA polymerase strategy, where the RNA polymerase first synthesizes either the digoxigenin or biotin labeled fragment of the ssRNA during IVTR, then is stalled after several cycles of biotin/DIG labeled UTP incorporation. The complexes are then purified to remove the labeled UTPs, and re-started in elongation to finalize the ssRNA strands, which are eventually annealed together (30). The stalled RNA polymerase strategy cannot benefit from the IVTR ssRNA amplification – ~10-fold typically – and has therefore a very limited yield. We believe that the ANS method we present here is simpler and more efficient to generate coilable dsRNA scaffolds.

### LNS versus ANS to fabricate NA hairpin scaffolds

We evaluated the LNS and the ANS strategies to generate NA hairpins with a long stem, i.e. hundreds of base pairs. Similarly to the linear coilable scaffolds, the batch of hairpin is separated in three fractions: the tethers opening as hairpins upon force increase (**Figure 5AB**), the tethers that do not open upon force increase, and junk nucleic acid that did not assemble into tethers during the fabrication. To quantify the fraction of functional hairpins in the batch, we first identified the NA tethers that show sudden tether length increase, i.e. hairpin opening, when the increasing force reaches a critical threshold (**Figure 5AB**). Second, we measured the fraction of hairpin tethers obtained per nanogram (ng) of NA construct per FoV to obtain the yield of the batch.

**Figure 5.**
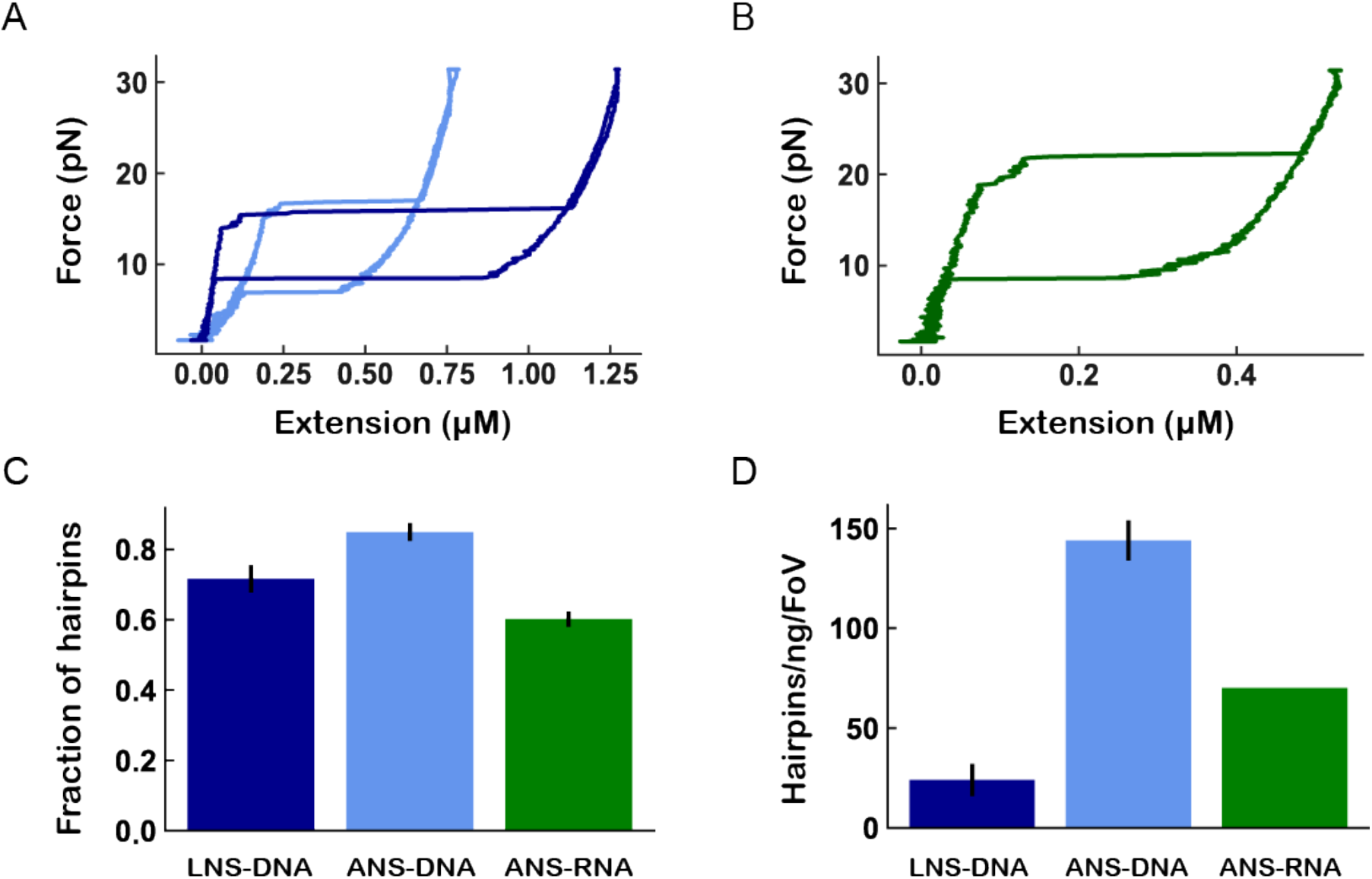
Comparative assessment of quality and yield for LNS and ANS methods. **(A)** Blue and dark blue traces are respectively the ANS and LNS DNA hairpins extension as a function of force, as described in **Figure 1B**. **(B)** Extension as a function of force for RNA hairpin. **(C)** Fraction of NA hairpins relative to the total number of tethers for DNA or RNA constructs assembled using LNS or ANS strategies. **(D)** Number of hairpin tethers obtained per nanogram of nucleic acid scaffold per field of view.

We observed that the fraction of functional hairpins in the LNS and ANS DNA hairpin samples, and the RNA hairpin sample were 0.72 ± 0.05, 0.85 ± 0.02 and 0.60 ± 0.02 (median ± 95% confidence interval), respectively (**Figure 5C, Supplementary Table 3**). Importantly, the number of functional hairpins per ng of used DNA was six times higher for ANS (144 hairpins in average per field of view) compared to LNS (24 hairpins). The total amounts of DNA in the batches were similar, i.e. 1535 ng for ANS and 1390 ng for LNS. These numbers clearly show that the ANS strategy produces more than 6-fold the amount of functional hairpins in one batch of manufactured scaffold sample than the LNS strategy. The main reason for the superiority of ANS stems from the fact that it contains less junk DNA, i.e. ANS uses more efficiently the initial ssDNA’s. Finally, the assembly of a DNA hairpin scaffold with the ANS strategy requires fewer steps and is faster than with the LNS method, and therefore less prone to human mistake.

Altogether, these advantages make the ANS strategy particularly attractive to generate long stem DNA hairpins. Also, it is well suited to produce long stem RNA hairpins for single molecule force spectroscopy experiments.

## Discussion

Scaffold fabrication is an essential prerequisite to study protein-nucleic acid interactions or mechanical properties of nucleic acids using force/torque spectroscopy techniques. Therefore, any laboratory performing such experiments has first to develop efficient and reliable protocols to generate these NA scaffolds. In the past few years, several research groups have published specific protocols related to one particular application (17,22–28). Though these studies have produced landmark articles on the investigated specific biomolecular processes, there has not been a comparative study that provides a clear overview of alternative methodologies used to prepare different types of DNA and RNA scaffolds. Here, we provide a clear overview of alternative methodologies used to prepare different types of DNA and RNA scaffolds. Importantly, we have developed a new strategy, named ANS, to generate linear coilable or long stem hairpin scaffolds, both made from either DNA or RNA. We have compared the ANS strategy to the most standard approach of preparing DNA scaffolds by ligating PCR products (LNS strategy). Using the ANS strategy, we are able to generate large amounts of either linear doublestranded or long stem hairpin NA with a high purity, an essential point for single molecule force and torque spectroscopy experiments. We show that ANS is especially well fitted to generate complex scaffolds that would otherwise require multiple ligation-purification steps, such as a DNA hairpin. However, we believe that for simpler scaffolds, e.g. linear dsDNA, the traditional LNS strategy may be better suited, as it requires just a single ligation-purification step, which only mildly affects both the final recovery yield and the coilable fraction. Concerning the fabrication of linear coilable dsRNA and long stem RNA hairpin scaffolds, we present here new protocols that provide both high yield and high quality linear coilable dsRNA and long stem hairpin, and therefore represent very useful alternatives to existing strategies (28,30). Long stem RNA hairpin will be particularly interesting to investigate kinetics RNA virus replicase, i.e. viral polymerase and helicase.

Finally, we believe that the present work will be useful not only to the single molecule force and torque spectroscopy community, but also to the single molecule fluorescence community, as complex NA scaffolds are also assembled to generate, e.g. labeled DNA for single molecule FRET study of bacterial transcription initiation (41).

## Data availability

The data of this study are available from the authors upon reasonable request.

## Acknowledgements

DD was supported by the Interdisciplinary Center for Clinical Research (IZKF) at the University Hospital of the University of Erlangen-Nuremberg. We would like to thank Eugen Ostrofet and Monika Spermann for helping with initial experiments and discussions, and Theo van Laar for assistance in the protocol design for the LNS strategy. We thank Anssi Malinen and Francesco Pedaci for careful reading of the manuscript.

## Author contributions

DD designed and supervised the research. FSP and DD designed the nucleic acid constructs. FSP made the nucleic acid constructs. MS performed the single molecule magnetic tweezers experiments, analyzed and plotted the data. FSP and DD wrote the article.

