## Supplementary Information for "High-yield fabrication of DNA and RNA scaffolds for single molecule force and torque spectroscopy experiments"

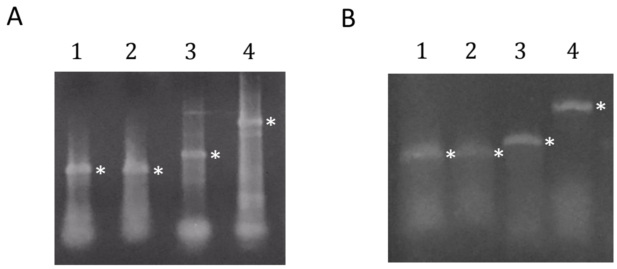


**Supplementary Figure 1. Separation of nicked from non-nicked DNA single strands in an agarose gel. (A)** Non-alkaline gel; **(B)** Alkaline gel. Purified bands are marked with an asterisk.

1) Biotin handle, 2) Digoxygenin handle, 3) Fragment S, 4) Fragment A.

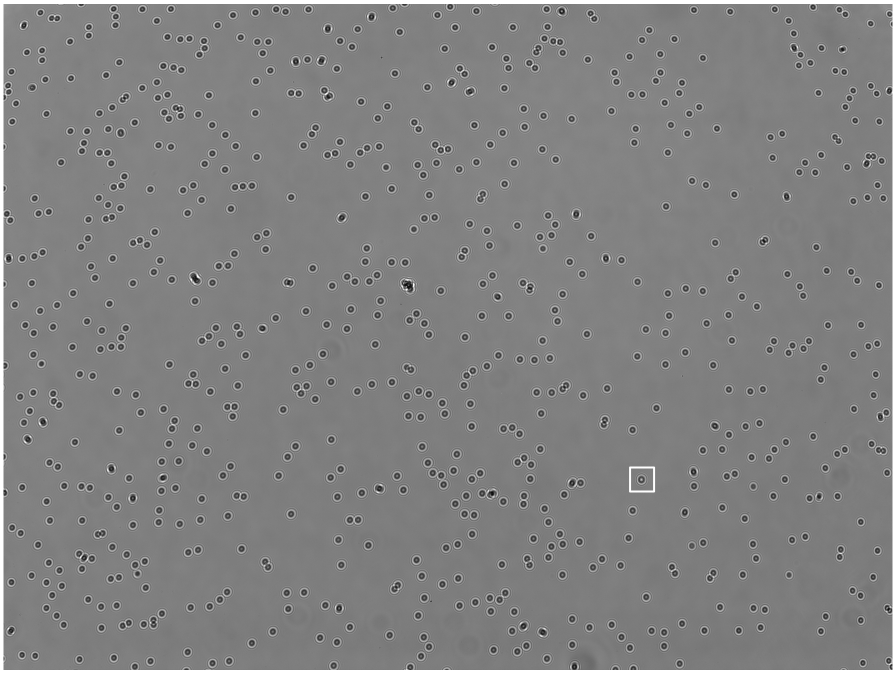

**Supplementary Figure 2.** Typical field of view from a high throughput magnetic tweezers experiment, where 1 μm diameter MyOne magnetic beads (dark circles) are tethered to the glass surface by a ~1.4 kbp coilable DNA construct. The polystyrene reference bead is highlighted by a white square.

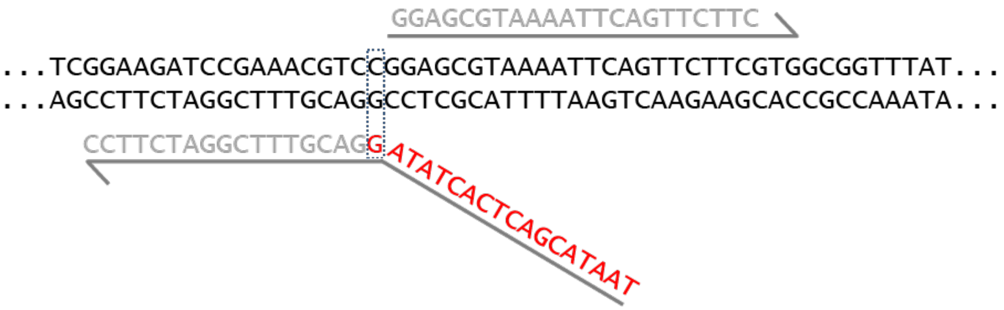


**Supplementary Figure 3.** Schematic of the primers (grey) design to insert the T7 RNAP promoter (red) in the sequence (black) to subsequently perform a successful ligation on RNA strands.

**Protocols to fabricate DNA and RNA scaffolds using the ANS strategy:**

Here we present the detailed protocols to generate DNA and RNA scaffolds for single-molecule force and torque spectroscopy experiments using the ANS strategy.

Materials

*Reagents*
Biotin-16-deoxyuridine triphosphate (dUTP) (1 mM; Roche)
Digoxigenin-11-dUTP, alkali stable (1 mM; Roche)
Plasmid DNA for amplification (**Supplementary Table 1**)*
Primers for amplification (**Supplementary Tables 1 and 2**)
QIAquick PCR Purification Kit (Qiagen) or
Monarch™ PCR & DNA Cleanup Kit (NEB)
Monarch™ DNA Gel Extraction Kit (NEB)
Restriction enzymes: BsaI , Nb.BbvCI, Nt.BbvCI (NEB)
T4 DNA ligase (10 U/μl) and 10× reaction buffer (NEB)
Taq PCR kit (NEB)
Phusion High-Fidelity PCR kit (NEB)
LA PCR™ Kit (Takara)
T4 DNA Polymerase, 3 U/μl (NEB)
HiScribe T7 High Yield RNA Synthesis kit (NEB)
Antarctic Phosphatase, 5 U/μl (NEB)
T4 Polynucleotide Kinase, 10 U/μl (NEB)
T4 RNA ligase 2, 10 U/μl (NEB)
Superase 20 U/μl (Invitrogen)
RNA Clean & Concentrator-5 kit (Zymo Research)
Zymoclean Gel RNA Recovery Kit (Zymo Research)
20x SSC (3 M Sodium Chloride and 0.3 M Sodium Citrate)
1 mM Sodium Citrate, pH 6.4 (The RNA Storage Solution, Invitrogen)
TAE (Tris-acetate-EDTA) buffer
Agarose (Sigma-Aldrich)
SyBr™Gold Nucleic Acid Gel Stain (Invitrogen)
RNA Gel Loading Dye (2X) (Invitrogen)

*Equipment*
Thermocycler
Microcentrifuge
Precision balance
Chamber for gel electrophoresis with power supply
Microwave oven

**For ANS DNA constructs, template DNA must contain all segments of the final construct in tandem and with evenly distributed BbvCI recognition sites. If sequence specificity is important for the study, the corresponding relevant parts of the sequence should not have BbvCI sites.*

Procedure

Set up two amplification reactions for each DNA fragment (or more, when noted), in 0.2 ml PCR tubes, keeping them on ice during the whole procedure.

***ANS linear coilable DNA***

1 To amplify DNA fragments for the linear coilable construct (**Figure 2B**), prepare reactions as follows:

| Reagent | BIO handle | DIG handle |
|---|---|---|
| 10x Standard Taq buffer | 5 µl | 5 µl |
| dNTPs (2.5 mM each) | 1 µl | 1 µl |
| primer P11 (10 µM) | 1 µl | - |
| primer P12 (10 µM) | 1 µl | - |
| primer P7 (10 µM) | - | 1 µl |
| primer P8 (10 µM) | - | 1 µl |
| Biotin-16-dUTP (1mM) | 2 µl | - |
| Digoxygenin-11-dUTP (1mM) | - | 2 µl |
| Template DNA (1 ng/µl) | 1 µl | 1 µl |
| Taq polymerase | 0.5 µl | 0.5 µl |
| Water | 38.5 µl | 38.5 µl |

2 Place the tubes in thermocycler and amplify DNA according to the following program:

| Cycle number | Denaturation | Annealing | Polymerization |
|---|---|---|---|
| 1 | 30 s at 95°C | | |
| 2–35 | 10 s at 95 °C | 15 s at 52 °C | 1 min 15 s at 68 °C |
| 36 | | | 5 min at 68 °C |

3 For the stem (S) and for fragment A, prepare four amplification reactions with the high fidelity Phusion polymerase, as follows:

| Reagent | S | A |
|---|---|---|
| 5X HF Buffer | 10 µl | 10 µl |
| dNTPs (2.5 mM each) | 1 µl | 1 µl |
| primer P9 (10 µM) | 1.5 µl | 1.5 µl |
| primer P10 (10 µM) | 1.5 µl | 1.5 µl |
| Template DNA (1 ng/µl) | 1 µl | 1 µl |
| Phusion polymerase | 0.5 µl | 0.5 µl |
| Water | 34.5 µl | 34.5 µl |

4 Place the tubes in thermocycler and amplify DNA according to the following program:

| Cycle number | Denaturation | Annealing | Polymerization |
|---|---|---|---|
| 1 | 30 s at 98°C | | |
| 2–35 | 10 s at 98 °C | 15 s at 52 °C | 1 min 15 s at 72 °C |
| 36 | | | 5 min at 72 °C |

5 Check fragment size in a 0.8% agarose gel. Pool replicate samples and purify using the QIAquick PCR Purification Kit. Adjust concentration of samples to 100 ng/µl.

6 Incubate BIO and DIG purified PCR products with T4 DNA Polymerase to remove 3' A overhangs left by the Taq polymerase. This step is critical to enable ligation of nicks after annealing of SSDNA strands. Prepare reactions as follows:

| **Reagent** | |
|---|---|
| Purified PCR product | 40 µl |
| 10x Cutsmart buffer | 5 µl |
| dNTPs (2.5 mM each) | 0.5 µl |
| T4 DNA polymerase | 1.3 µl |
| Water | 4.2 µl |

Incubate reactions for 15 min at 12°C. Place on ice.

7 Purify reactions using the Monarch™ PCR & DNA Cleanup Kit. Adjust concentration to 50 ng/µl.

8 Set up reactions to nick one strand of the DNA:

| **Reagent** | **BIO handle** | **DIG handle** | **S** | **A** |
|---|---|---|---|---|
| Template DNA (PCR) | 30 µl | 30 µl | 30 µl | 30 µl |
| 10x Cutsmart buffer | 4 µl | 4 µl | 4 µl | 4 µl |
| Nb.BbvCI | 1 µl | 1 µl | 1 µl | - |
| Nt.BbvCI | - | - | - | 1 µl |
| Water | 5 µl | 5 µl | 5 µl | 5 µl |

Incubate at 37°C for 2 h. Purify samples using the Monarch™ PCR & DNA Cleanup Kit and elute in 10µl buffer to concentrate samples.

9 Prepare a 1.2 % agarose gel in TAE buffer. Add 10 µl of RNA Gel Loading Dye (denaturing) to each sample and heat at 75°C for 5 min. Place tubes on ice or at 4°C immediately. Load samples onto gel and run electrophoresis at 5 V/cm for two hours. Stain gel with 1x SyBr™Gold dye for 15 min and remove non-incorporated dye by immersing the gel in TAE buffer for aproximately 30 min.

10 Cut gel slices containing the desired fragments (see **Supplementary Figure 1B** for example) and purify them using the Monarch™ DNA Gel Extraction Kit. Follow manufacturer's recommendation for ssDNA purification and use 7 volumes of gel dissolving buffer to the agarose slice (e.g., 700 µl buffer for 100 mg agarose).

11 Mix equimolar amounts (ca. 0.5 – 1 pmol) of each single-stranded fragment in a 0.2 ml tube and add NaCl to a final concentration of 50 mM. Incubate samples in a pre-programmed thermocycler with an initial cycle of 65°C for one hour and then cool down by 1.2°C/5 minutes, until it reaches 25°C.

12 Add 1/10 vol of 10x T4 DNA ligase buffer to the reaction and 1.5 µl of DNA Ligase and incubate 5 min at room temperature or overnight at 16°C.

13 Aliquot and store samples at -80 °C.

***ANS DNA hairpin***

1 To amplify DNA fragments for the hairpin construct (**Figure 3B**), prepare two reactions for each DNA fragment as follows:

| **Reagent** | **BIO handle** | **DIG handle** | **C1** | **C2** |
|---|---|---|---|---|
| 10x Standard Taq buffer | 5 µl | 5 µl | 5 µl | 5 µl |
| dNTPs (2.5 mM each) | 1 µl | 1 µl | 1 µl | 1 µl |
| primer P17 (10 µM) | 1 µl | - | - | - |
| primer P18 (10 µM) | 1 µl | - | 1 µl | - |
| primer P19 (10 µM) | - | 1 µl | - | 1 µl |
| primer P20 (10 µM) | - | 1 µl | - | - |
| primer P21 (10 µM) | - | - | 1 µl | - |
| primer P22 (10 µM) | - | - | - | 1 µl |
| Biotin-16-dUTP (1mM) | 2 µl | - | - | - |
| Digoxygenin-11-dUTP (1mM) | - | 2 µl | - | - |
| Template DNA (1 ng/µl) | 1 µl | 1 µl | 1 µl | 1 µl |
| Taq polymerase | 0.5 µl | 0.5 µl | 0.5 µl | 0.5 µl |
| Water | 38.5 µl | 38.5 µl | 40.5 µl | 40.5 µl |

2 Place the tubes in thermocycler and amplify DNA according to the following program:

| **Cycle number** | **Denaturation** | **Annealing** | **Polymerization** |
|---|---|---|---|
| 1 | 30 s at 95°C | | |
| 2–35 | 10 s at 95 °C | 15 s at 52 °C | 1 min at 68 °C |
| 36 | | | 5 min at 68 °C |

3 Amplify the S-SL fragment, which contains the complementary inverted repeat, using the LA Taq polymerase.

| **Reagent** | **S-SL** |
|---|---|
| 2X GC Buffer I | 25 µl |
| dNTPs (2.5 mM each) | 1 µl |
| primer P23 (10 µM) | 1 µl |
| primer P24 (10 µM) | 1 µl |
| Template DNA (1 ng/µl) | 1 µl |
| LA Taq polymerase | 0.5 µl |
| Water | 20.5 µl |

4 Place the tubes in thermocycler and amplify DNA according to the following program:

| **Cycle number** | **Denaturation** | **Annealing** | **Polymerization** |
|---|---|---|---|
| 1 | 2 min at 94°C | | |
| 2–30 | 20 s at 94 °C | 15 s at 52 °C | 2 min at 68 °C |
| 36 | | | 5 min at 68 °C |

5 Check fragment size in a 0.8% agarose gel. Pool replicate samples and purify using the QIAquick PCR Purification Kit. Due to the presence of the complementary inverted

repeat, the S-SL fragment should be purified from an agarose gel, since multiple bands are possible. Adjust concentration to 50 ng/µl.

6 Nick the purified PCR fragments using the appropriate endonucleases, following the schematic below:

| **Reagent** | **BIO handle** | **DIG handle** | **C1** | **C2** | **S-SL** |
|---|---|---|---|---|---|
| Template DNA (PCR) | 40 µl | 40 µl | 40 µl | 40 µl | 40 µl |
| 10x Cutsmart buffer | 5 µl | 5 µl | 5 µl | 5 µl | 5 µl |
| Nb.BbvCI | 1 µl | 1 µl | - | - | 1 µl |
| Nt.BbvCI | - | - | 1 µl | 1 µl | - |
| Water | 4 µl | 4 µl | 4 µl | 4 µl | 4 µl |

Incubate reactions at 37°C for 2 h. Heat samples at 80°C for 20 min to inactivate enzymes and detach ~80 nt oligonucleotides resulting from the multiple nicking of one strand.

7 Submit digestion products to agarose gel electrophoresis under alkaline conditions (see **Materials and Methods**).

8 Cut gel slices containing the desired fragments (see **Supplementary Figure 1B** for example) and purify them using the Monarch™ DNA Gel Extraction Kit. Follow manufacturer's recommendation for ssDNA purification and use 7 volumes of gel dissolving buffer to the agarose slice (e.g., 700 µl buffer for 100 mg agarose).

9 Mix equimolar amounts (ca. 1 – 2 pmol) of each single-stranded fragment in a 0.2 ml tube and add NaCl to a final concentration of 50 mM. Incubate samples in a pre-programmed thermocycler with an initial cycle of 65°C for one hour and then cool down by 1.2°C/5 minutes, until it reaches 25°C.

10 Aliquot and store samples at -80 °C.

***ANS linear coilable RNA***

1 Amplify DNA fragments via PCR to obtain the templates for in vitro transcription (IVTR). For the linear coilable RNA construct, prepare reactions as follows:

| Reagent | BIO handle | DIG handle | SPCD | AB |
|---|---|---|---|---|
| 5X HF Buffer | 10 µl | 10 µl | 10 µl | 10 µl |
| dNTPs (2.5 mM each) | 1 µl | 1 µl | - | - |
| primer P33 (10 µM) | 1.5 µl | - | - | - |
| primer P34 (10 µM) | 1.5 µl | - | - | - |
| primer P37 (10 µM) | - | 1.5 µl | - | - |
| primer P38 (10 µM) | - | 1.5 µl | - | - |
| primer P35 (10 µM) | - | - | 1.5 µl | - |
| primer P36 (10 µM) | - | - | 1.5 µl | - |
| primer P34 (10 µM) | - | - | - | 1.5 µl |
| primer P39 (10 µM) | - | - | - | 1.5 µl |
| Template DNA (1 ng/µl) | 1 µl | 1 µl | - | - |
| Phusion polymerase | 0.5 µl | 0.5 µl | - | - |
| Water | 34.5 µl | 34.5 µl | - | - |

2 Place the tubes in thermocycler and amplify DNA according to the following program:

| Cycle number | Denaturation | Annealing | Polymerization |
|---|---|---|---|
| 1 | 30 s at 98°C | | |
| 2–35 | 10 s at 98 °C | 15 s at 52 °C | 1 min 15 s at 72 °C |
| 36 | | | 5 min at 72 °C |

3 Purify samples using Monarch PCR & DNA Cleanup Kit and elute in 20 µl to concentrate samples.

4 Set up in vitro transcription reactions using the HiScribe T7 High Yield RNA Synthesis kit to obtain RNA. Use ~500 ng of template DNA per 20 µl reaction. Proceed as follows:

| Reagent | BIO handle | DIG handle | SPCD | AB |
|---|---|---|---|---|
| 10 x reaction buffer | 1.5 µl | 1.5 µl | 2 µl | 2 µl |
| ATP (100 mM) | 1.5 µl | 1.5 µl | 2 µl | 2 µl |
| CTP (100 mM) | 1.5 µl | 1.5 µl | 2 µl | 2 µl |
| GTP (100 mM) | 1.5 µl | 1.5 µl | 2 µl | 2 µl |
| UTP (100 mM) | 1 µl | 1 µl | 2 µl | 2 µl |
| Biotin-16-UTP (10 mM) | 5 µl | - | - | - |
| Digoxygenin-11-UTP (10 mM) | - | 5 µl | - | - |
| Template DNA (1 ng/µl) | up to 6 µl | 6 µl | 7.5 µl | 7.5 µl |
| Superase | 0.5 µl | 0.5 µl | 0.5 µl | 0.5 µl |
| T7 Polymerase mix | 1.5 µl | 1.5 µl | 2 µl | 2 µl |
| Nuclease-free water | q.s.p 20 µl | 20 µl | 20 µl | 20 µl |

Incubate samples for 2h at 37°C (preferably in the Thermocycler to avoid evaporation).

5 Purify reactions with the RNA Clean & Concentrator-5 kit. Adjust concentration to 600 ng/µl and treat RNA with Antarctic Phosphatase to remove 5'-triphosphates, as follows:

| Reagent | BIO handle | DIG handle | SPCD | AB |
|---|---|---|---|---|
| 10 x AP reaction buffer | 3 µl | 3 µl | 3 µl | 3 µl |
| RNA | 8 µl | 8 µl | 8 µl | 8 µl |
| Antarctic Phosphatase | 1 µl | 1 µl | 1 µl | 1 µl |
| Nuclease-free water | 18 µl | 18 µl | 18 µl | 18 µl |

Incubate samples 1h at 37°C.

6 Purify reactions with the RNA Clean & Concentrator-5 kit. Add one phosphate group (required for ligation) to the 5'-end of the RNA strands using the T4 Protein Kinase, as described below:

| Reagent | BIO handle | DIG handle | SPCD | AB |
|---|---|---|---|---|
| 10 x T4 PNK reaction buffer | 3 µl | 3 µl | 3 µl | 3 µl |
| RNA | 8 µl | 8 µl | 8 µl | 8 µl |
| T4 Protein kinase | 1 µl | 1 µl | 1 µl | 1 µl |
| Nuclease-free water | 18 µl | 18 µl | 18 µl | 18 µl |

7 Purify reactions with the RNA Clean & Concentrator-5 kit. Elute in 15 µl.

8 Prepare annealing reactions by mixing equimolar amounts (ca. 1 – 2 pmol) of each RNA sample in a 0.2 ml tube. Add 2.5 µl of 20x SSC and 1 mM sodium citrate buffer to a final reaction volume of 100 µl. Incubate samples in a pre-programmed thermocycler with an initial cycle of 65°C for one hour and then cool down by 1.2°C/5 minutes, until it reaches 25°C.

9 Concentrate samples using the RNA Clean & Concentrator-5 kit and elute in 17µl nuclease-free water. Prepare a ligation reaction using 17 µl dsRNA, 2 µl T4 ligase 2 buffer and 1 µl T4 RNA ligase 2. Incubate 1 h at 37°C.

10 (optional) Purify with the RNA Clean & Concentrator-5 kit and elute in 15 µl of 1 mM Sodium Citrate, pH 6.4. Dilute and aliquot, if necessary, and store at -80°C.

*ANS RNA hairpin*

1 Amplify DNA fragments via PCR to obtain the templates for in vitro transcription (IVTR). For the RNA hairpin construct, prepare reactions as follows:

| **Reagent** | **BIO** | **DIG** | **RS1** | **RS2** | **loop** | **C1** | **C2** |
|---|---|---|---|---|---|---|---|
| 5X HF Buffer | 10 µl | 10 µl | 10 µl | 10 µl | 10 µl | 10 µl | 10 µl |
| dNTPs (2.5 mM each) | 1 µl | 1 µl | | 1 µl | 1 µl | 1 µl | 1 µl |
| primer P42 (10 µM) | 1.5 µl | - | - | - | - | - | - |
| primer P43 (10 µM) | 1.5 µl | - | - | - | - | - | - |
| primer P40 (10 µM) | - | 1.5 µl | - | - | - | - | - |
| primer P20 (10 µM) | - | 1.5 µl | - | - | - | - | - |
| primer P24 (10 µM) | - | - | 1.5 µl | - | - | - | - |
| primer P49 (10 µM) | - | - | 1.5 µl | - | - | - | - |
| primer P41 (10 µM) | - | - | - | 1.5 µl | - | - | - |
| primer P50 (10 µM) | - | - | - | 1.5 µl | - | - | - |
| primer P47 (10 µM) | - | - | - | - | 1.5 µl | - | - |
| primer P48 (10 µM) | - | - | - | - | 1.5 µl | - | - |
| primer P44 (10 µM) | - | - | - | - | - | 1.5 µl | - |
| primer P45 (10 µM) | - | - | - | - | - | 1.5 µl | - |
| primer P46 (10 µM) | - | - | - | - | - | - | 1.5 µl |
| primer P19 (10 µM) | - | - | - | - | - | - | 1.5 µl |
| Template DNA (1 ng/µl) | 1 µl | 1 µl | 1 µl | 1 µl | 1 µl | 1 µl | 1 µl |
| Phusion polymerase | 0.5 µl | 0.5 µl | 0.5 µl | 0.5 µl | 0.5 µl | 0.5 µl | 0.5 µl |
| Water | 34.5 µl | 34.5 µl | 34.5 µl | 34.5 µl | 34.5 µl | 34.5 µl | 34.5 µl |

2 Place the tubes in thermocycler and amplify DNA according to the following program:

| **Cycle number** | **Denaturation** | **Annealing** | **Polymerization** |
|---|---|---|---|
| 1 | 30 s at 98°C | | |
| 2–35 | 10 s at 98 °C | 15 s at 52 °C | 1 min at 72 °C |
| 36 | | | 5 min at 72 °C |

3 Purify samples using Monarch PCR & DNA Cleanup Kit and elute in 20 µl to concentrate samples.

4 Set up in vitro transcription reactions using the HiScribe T7 High Yield RNA Synthesis kit to obtain RNA. Use ~500 ng of template DNA per 20 µl reaction. Proceed as follows:

| **Reagent** | **BIO C2** | **DIG** | **RS1** | **RS2** | **loop** | **C1** | |
|---|---|---|---|---|---|---|---|
| 10 x reaction buffer | 1.5 µl | 1.5 µl | 2 µl | 2 µl | 2 µl | 2 µl | 2 µl |
| ATP (100 mM) | 1.5 µl | 1.5 µl | 2 µl | 2 µl | 2 µl | 2 µl | 2 µl |
| CTP (100 mM) | 1.5 µl | 1.5 µl | 2 µl | 2 µl | 2 µl | 2 µl | 2 µl |
| GTP (100 mM) | 1.5 µl | 1.5 µl | 2 µl | 2 µl | 2 µl | 2 µl | 2 µl |

| | | | | | | | |
|---|---|---|---|---|---|---|---|
| UTP (100 mM) | 1 µl | 1 µl | 2 µl | 2 µl | 2 µl | 2 µl | 2 µl |
| Biotin-16-UTP (10 mM) | 5 µl | - | - | - | - | - | - |
| Dig-11-UTP (10 mM) | - | 5 µl | - | - | - | - | - |
| Template DNA (1 ng/µl) | 6 µl | 6 µl | 7.5 µl | 7.5 µl | 7.5 µl | 7.5 µl | 7.5 µl |
| Superase | 0.5 µl | 0.5 µl | 0.5 µl | 0.5 µl | 0.5 µl | 0.5 µl | 0.5 µl |
| T7 Polymerase mix | 1.5 µl | 1.5 µl | 2 µl | 2 µl | 2 µl | 2 µl | 2 µl |

Incubate samples for 2h at 37°C (preferably in the Thermocycler to avoid evaporation).

5 Purify reactions with the RNA Clean & Concentrator-5 kit. Adjust concentration to 600 ng/µl and treat RNAs that will have to be ligated (BIO, RS1, RS2 and loop) with Antarctic Phosphatase to remove 5'-triphosphates, as follows:

| **Reagent** | **BIO** | **RS1** | **RS2** | **loop** |
|---|---|---|---|---|
| 10 x AP reaction buffer | 3 µl | 3 µl | 3 µl | 3 µl |
| RNA | 8 µl | 8 µl | 8 µl | 8 µl |
| Antarctic Phosphatase | 1 µl | 1 µl | 1 µl | 1 µl |
| Nuclease-free water | 18 µl | 18 µl | 18 µl | 18 µl |

Incubate samples 1h at 37°C.

6 Purify reactions with the RNA Clean & Concentrator-5 kit. Add one phosphate group (required for ligation) to the 5'-end of the RNA strands using the T4 Protein Kinase, as described below:

| **Reagent** | **BIO** | **RS1** | **RS2** | **loop** |
|---|---|---|---|---|
| 10 x T4 PNK reaction buffer | 3 µl | 3 µl | 3 µl | 3 µl |
| RNA | 8 µl | 8 µl | 8 µl | 8 µl |
| T4 Protein kinase | 1 µl | 1 µl | 1 µl | 1 µl |
| Nuclease-free water | 18 µl | 18 µl | 18 µl | 18 µl |

7 Purify reactions with the RNA Clean & Concentrator-5 kit. Elute in 15 µl.

8 Prepare annealing reactions by mixing equimolar amounts (ca. 1 – 2 pmol) of each RNA sample in a 0.2 ml tube. Add 2.5 µl of 20x SSC and 1 mM sodium citrate buffer to a final reaction volume of 100 µl. Incubate samples in a pre-programmed thermocycler with an initial cycle of 65°C for one hour and then cool down by 1.2°C/5 minutes, until it reaches 25°C.

9 Concentrate samples using the RNA Clean & Concentrator-5 kit and elute in 17µl nuclease-free water. Prepare a ligation reaction using 17 µl dsRNA, 2 µl T4 ligase 2 buffer and 1 µl T4 RNA ligase 2. Incubate 1 h at 37°C.

10 (optional) Purify with the RNA Clean & Concentrator-5 kit and elute in 15 µl of 1 mM Sodium Citrate, pH 6.4. Dilute and aliquot, if necessary, and store at -80°C.

**Supplementary Table 1. List of constructs and primers used in this study.** For primer sequences refer to **Supplementary Table 2**.

| Strategy | Scaffold type | Template DNA | Fragment name | Primers for PCR | Fragment size |
|---|---|---|---|---|---|
| LNS | Linear DNA | pMTT2 | stem | P1, P2 | 2070 bp |
| | | λ DNA | BIO handle | P3, P4 | 421 bp |
| | | | DIG handle | P5, P6 | 421 bp |
| | Hairpin DNA | pMTM | stem | P13, P14 | 1043 bp |
| | | λ DNA | BIO handle | P3, P4 | 421 bp |
| | | | DIG handle | P5, P6 | 421 bp |
| | | | spacer | P15, P16 | 519 bp |
| ANS | Linear DNA | pP1T1T2 | DIG handle | P7, P8 | 900 bp |
| | | | S (stem) | P9, P10 | 1400 bp |
| | | | BIO handle | P11, P12 | 955 bp |
| | | | A | P7, P12 | 3255 bp |
| | Hairpin DNA | pMTM2 | BIO handle | P17, P18 | 404 bp |
| | | | DIG handle | P19, P20 | 343 bp |
| | | | S-SL | P23, P24 | 1974 bp |
| | | | C1 | P21, P18 | 847 bp |
| | | | C2 | P19, P22 | 856 bp |
| | Linear RNA nc | pBB10* | BIO handle | P25, P26 | 452 bp |
| | | | DIG handle | P27, P28 | 513 bp |
| | | | SP | P29, P30 | 354 bp |
| | | | CD | P31, P32 | 2820 bp |
| | | | AB | P33, P34 | 4144 bp |
| | Linear RNA c | pBB10* | BIO handle | P33, P34 | 447 bp |
| | | | DIG handle | P37, P38 | 515 bp |
| | | | SPCD | P35, P36 | 3173 bp |
| | | | AB | P34, P39 | 4135 bp |
| | Hairpin RNA | pMTM2 | BIO handle | P42, P43 | 404 bp |
| | | | DIG handle | P40, P20 | 343 bp |
| | | | RS1 | P24, P49 | 870 bp |
| | | | RS2 | P41, P50 | 1002 bp |
| | | | loop | P47, P48 | 103 bp |
| | | | C1 | P44, P45 | 822 bp |
| | | | C2 | P46, P19 | 856 bp |

Abbreviations: nc=non-coilable, c=coilable; *(Wang *et al.*, J. Bacteriol. 2006)

**Supplementary Table 2. Primer list.**

| # | name | sequence 5'-3' | restriction site | modification |
|---|---|---|---|---|
| P1 | MTT_STEM_F | AAAATAGGAGAGACCCTTTGGATCCCGTCATTGCG | BsaI | |
| P2 | MTT_STEM_R | AAAAGGTCTCTGCAAAGTAAAGCTTAGTGTCACGC | BsaI | |
| P3 | BIOh_F | GGAACCAAAGGATATTCAGACGCG | | |
| P4 | BIOh_R | AAAAGGTCTCATTGCGGATCCCGTGATGACCTC | BsaI | |
| P5 | DIGh_F | GGATCCCGTGATGACCTCATTA | | |
| P6 | DIGh_R | AAAACCTAAGAGACCGGAACCAAAGGATATTCAGACG | BsaI | |
| P7 | PTT_DIGh_F | CCTCACTTCTGCTATTTCGCAGG | | |
| P8 | PTT_DIGh_R | CAGCTGAGGACTCCAAACG | | |
| P9 | PTT_S_F | AACAAATCTATTATACCAATCGGCT | | 5'-P |
| P10 | PTT_S_R | CGATGGTGGTAGCGGC | | |
| P11 | PTT_BIOh_F | ATTAATACAACGAACGGTGATGTTG | | 5'-P |
| P12 | PTT_BIOh_R | GGAACCAAAGGATATTCAGACG | | |
| P13 | MTMhp_fw | CGACATACGGTTGAGACCG | | |
| P14 | MTMhp_rev | GCGCGGTCTCGGTATTATC | | |
| P15 | SPhp_F | AAAAGGTCTCATTGAAACCGGAGTGATGTCGCG | BsaI | |
| P16 | SPhp_R | AAAAGGTCTCTTAGGCTTCGCCAGACGGCAT | BsaI | |
| P17 | Ann_BIOh_F | GCAGGCAAGTCCGATTTTTTG | | |
| P18 | Ann_BIOh_R | GGAACCAAAGGATATTCAGACG | | |
| P19 | Ann_DIGh_F | CCTCACTTCTGCTATTTCGC | | |
| P20 | Ann_DIGh_R | GCGACTTAGCTGAGGCC | | |
| P21 | Ann_C1_F | CCTCAGCCAACTCGGTCG | | |
| P22 | Ann_C2_R | TGCTGAGGAACCGGAGTG | | |
| P23 | Ann_stem_F | TCTTCGCCAGACGGCATTTA | | |
| P24 | Ann_stem_R | GGTGCCCACAGAACGTC | | |
| P25 | LinRNA_BIO_F | AAGATTAGCGGATCCTACCTGAC | | |
| P26 | LinRNA_BIO_R | **TAATACGACTCACTATAG**GAACGGCTTGATATCCACTTTACG | | |
| P27 | LinRNA_DIG_F | AGCGTAAAATTCAGTTCTTCGTGGCG | | |
| P28 | LinRNA_DIG_R | **TAATACGACTCACTATAG**GGGTTAACCTCAACTTCCATTTCC | | |
| P29 | LinRNA_SP_F | TGCCATTCAGGGACTGCCGATGTCGGTGCAGCCG | | |
| P30 | LinRNA_SP_R | **TAATACGACTCACTATAG**GAGCGCCGCTTCCATGTCCTGGAACGCT | | |
| P31 | LinRNA_CD_F | GGGAAAAAAAAAAAACCGTATGACGCTGGAAG | | |
| P32 | LinRNA_CD_R | **TAATACGACTCACTATAG**GCCGGACGTTTCGGATCTTCCGACATGCGC | | |
| P33 | LinRNA_AB_F | **TAATACGACTCACTATAG**GAAGATTAGCGGATCCTACCTGAC | | |
| P34 | LinRNA_AB_R | GGTTAACCTCAACTTCCATTTCC | | |
| P33 | CoilRNA_BIO_F | GGATCCTACCTGACGCTTTT | | |
| P34 | CoilRNA_BIO_R | **TAATACGACTCACTATAG**GCAAACGGCTTGATATCC | | |

| | | | |
|---|---|---|---|
| P35 | CoilRNA_SP_F | ATTCAGGGACTGCCGATG | |
| P36 | CoilRNA_CD_R | **TAATACGACTCACTATAG**GACGTTTCGGATCTTCC | |
| P37 | CoilRNA_DIG_F | GGAGCGTAAAATTCAGTTCTTC | |
| P38 | CoilRNA_DIG_R | **TAATACGACTCACTATAG**GTTAACCTCAACTTCCATTTCC | |
| P39 | CoilRNA_AB_F | **TAATACGACTCACTATAG**GATCCTACCTGACGCTTTT | |
| P40 | hpRNA_DIGh_F | **TAATACGACTCACTATAG**GCCTCACTTCTGCTATTTCGC | |
| P41 | hpRNA_Stem_F | **TAATACGACTCACTATAG**GTCTTCGCCAGACGGCATTTA | |
| P42 | hpRNA_BIO_F | **TAATACGACTCACTATAG**GGCAGGCAAGTCCGATTTTTTG | |
| P43 | hpRNA_BIO_R | GGAACCAAAGGATATTCAGACG | |
| P44 | hpRNA_C1_F | **TAATACGACTCACTATAG**GGGAACCAAAGGATATTCAGACG | |
| P45 | hpRNA_C1_R | AACAAGAAACTTCCTTGGCTG | |
| P46 | hpRNA_C2_F | **TAATACGACTCACTATAG**GTGCTGAGGAACCGGAGTG | |
| P47 | RNAloop_fw | **TAATACGACTCACTATAG**GGTAGTGATTAACATTCGACAGCATGCGCAC | |
| P48 | RNAloop_rev | CAGGATCACGTTACCGCC | |
| P49 | hp_RS1_dirBIO | **TAATACGACTCACTATAG**GAACGGCGCCTATGACG | |
| P50 | hp_RS2_dirDIG | TGGATCCGTGGGCGC | |
| O1 | oligo1 | GCAAAAGTCATTCTGAGAATAGTGTATGCGGCGACCGAGTTG | 5'-P |
| O2 | oligo2 | ACCGCAGTACAATCTGCTCTGATGCCGCATGTCTGCACGCTATGTAGAACCAACTCGGTCGCCGCATACACTATTCTCAGAATGACTT | 5'-P |
| O3 | oligo3 | TCAATAGATGTGCTGCCCCTCAGTCCGTTGATACCTACTTGCT | |
| O4 | oligo4 | TCAAAGCAAGTAGGTATCAACGGACTGAGGGGCAGCACATCTATTGATTTTTTTTTTTTTTTTGTTCTACATAGCGTGCAGACATGCGGCATCAGAGCAGATTGTACTG | 5'-P |
| O5 | oligo5 | ATACAGCTCGCCGAGGCGAGAAATCGCCTCGGCGAGCT | 5'-P |

Abbreviations: modif=modification, P=Phosphate

Obs.: T7 promoter recognition sequence is shown in bold and starting site for the T7 polymerase is underlined.

**Supplementary Table 3. Magnetic tweezers experimental data obtained with LNS and ANS scaffolds. Data of two independently fabricated batches for each type of construct are presented. Median and mean ratios of the number of coilable tethers (for linear constructs) or hairpins versus the total numbers of tethers in the analyzed fields of view are shown (GT/TT).**

| Strategy | Scaffold type | Exp. | GT/ng/FoV | avgGT/ng /FoV | Median GT/TT | Mean GT/TT | ng produced /batch |
|---|---|---|---|---|---|---|---|
| LNS | Linear DNA | 1 | 46 (5)* | 51 | 0.32 ± 0.05 | 0.35 ± 0.05 | 407 |
| | | 2 | 57 (4) | | 0.39 ± 0.04 | | 309 |
| | Hairpin DNA | 1 | 16 (5) | 24 | 0.73 ± 0.04 | 0.72 ± 0.04 | 609 |
| | | 2 | 32 (4) | | 0.71 ± 0.04 | | 2170 |
| ANS | Linear DNA | 1 | 126 (4) | 142 | 0.39 ± 0.06 | 0.38 ± 0.05 | 160 |
| | | 2 | 157 (5) | | 0.36 ± 0.06 | | 420 |
| | Hairpin DNA | 1 | 154 (5) | 144 | 0.86 ± 0.02 | 0.85 ± 0.03 | 2080 |
| | | 2 | 134 (4) | | 0.84 ± 0.03 | | 990 |
| | Linear RNA nc | 1 | 129 (3) | 129 | 0.82 ± 0.03 | 0.82 ± 0.03 | 540 |
| | Linear RNA c | 1 | 393 (5) | 425 | 0.29 ± 0.03 | 0.26 ± 0.03 | 2200 |
| | | 2 | 457 (3) | | 0.23 ± 0.04 | | 2760 |
| | Hairpin RNA | 1 | 70 (10) | 70 | 0.60 ± 0.02 | 0.60 ± 0.02 | 1176 |

GT=good tethers (coilables or hairpins), TT=total number of tethers, FoV=field of view, ng=nanogram, avg=average

*between brackets: number of FoV analyzed
